# The polybasic cleavage site in the SARS-CoV-2 spike modulates viral sensitivity to Type I IFN and IFITM2

**DOI:** 10.1101/2020.12.19.423592

**Authors:** Helena Winstone, Maria Jose Lista, Alisha Reid, Suzanne Pickering, Katie J Doores, Chad Swanson, Stuart J D Neil

**Affiliations:** Department of Infectious Diseases, School of Immunology and Microbial Sciences, King’s College London, London, United Kingdom

**Author notes:** Corresponding author: Stuart Neil. These authors contributed equally to this work (Helena Winstone, Maria Jose Lista).

## Abstract

The cellular entry of severe acute respiratory syndrome-associated coronaviruses types 1 and 2 (SARS-CoV-1 and -2) requires sequential protease processing of the viral spike glycoprotein (S). The presence of a polybasic cleavage site in SARS-CoV-2 S at the S1/S2 boundary has been suggested to be a factor in the increased transmissibility of SARS-CoV-2 compared to SARS-CoV-1 by facilitating maturation of the S precursor by furin-like proteases in the producer cells rather than endosomal cathepsins in the target. We investigate the relevance of the polybasic cleavage site in the route of entry of SARS-CoV-2 and the consequences this has for sensitivity to interferons, and more specifically, the IFN-induced transmembrane (IFITM) protein family that inhibit entry of diverse enveloped viruses. We found that SARS-CoV-2 is restricted predominantly by IFITM2 and the degree of this restriction is governed by route of viral entry. Removal of the cleavage site in the spike protein renders SARS-CoV-2 entry highly pH- and cathepsin-dependent in late endosomes where, like SARS-CoV-1 S, it is more sensitive to IFITM2 restriction. Furthermore, we find that potent inhibition of SARS-CoV-2 replication by type I but not type II IFNs is alleviated by targeted depletion of IFITM2 expression. We propose that the polybasic cleavage site allows SARS-CoV-2 to mediate viral entry in a pH-independent manner, in part to mitigate against IFITM-mediated restriction and promote replication and transmission. This suggests therapeutic strategies that target furin-mediated cleavage of SARS-CoV-2 S may reduce viral replication through the activity of type I IFNs.

**IMPORTANCE:** The furin cleavage site in the S protein is a distinguishing feature of SARS-CoV-2 and has been proposed to be a determinant for the higher transmissibility between individuals compared to SARS-CoV-1. One explanation for this is that it permits more efficient activation of fusion at or near the cell surface rather than requiring processing in the endosome of the target cell. Here we show that SARS-CoV-2 is inhibited by antiviral membrane protein IFITM2, and that the sensitivity is exacerbated by deletion of the furin cleavage site which restricts viral entry to low pH compartments. Furthermore, we find that IFITM2 is a significant effector of the antiviral activity of type I interferons against SARS-CoV-2 replication. We suggest one role of the furin cleavage site is to reduce SARS-CoV-2 sensitivity to innate immune restriction, and thus may represent a potential therapeutic target for COVID-19 treatment development.

## INTRODUCTION

Severe acute respiratory syndrome coronavirus 2 (SARS-CoV-2) is a novel coronavirus that was identified in early 2020(Wu et al., 2020). Entry of SARS-CoV-2 into the cell is initiated by the spike glycoprotein binding to its receptor, angiotensin-converting enzyme 2 (ACE2) (Hoffmann, Kleine-Weber, Schroeder, et al., 2020). Spike is a type I transmembrane protein that is synthesised as a polyprotein precursor and requires two steps of proteolytic cleavage at the S1/S2 boundary and at the S2’ site in order to mediate fusion of the viral and cell membranes. Due to the insertion of four amino acids at the S1/S2 boundary of SARS-CoV-2 spike, with the sequence _681_**PRRA**|RSV_687_, SARS-CoV-2 spike contains a canonical furin-like protease cleavage site (Hoffmann, Kleine-Weber, Schroeder, et al., 2020). This allows the SARS-CoV-2 spike to be cleaved by furin-like proteases intracellularly prior to virion release. TMPRSS2 on the target cell surface and cathepsins B and L in endosomes may cleave the S2’ site and activate the fusion machinery depending on the relative availability of these enzymes. The presence of this site has been suggested to be important for determining viral tropism and transmission of SARS-CoV-2 (Andersen, Rambaut, Lipkin, Holmes, & Garry, 2020; Hoffmann, Kleine-Weber, & Pöhlmann, 2020; Peacock et al., 2020). However, the necessity for this site is cell-type dependent. It has been shown that this site can be lost after several passages in TMPRSS2-negative Vero-E6 cells (Davidson et al., 2020). Nevertheless, similar mutations have only been found rarely in a small number of patients (Liu et al., 2020; Sasaki et al., 2020). This suggests a selective pressure to conserve the polybasic cleavage site for *in vivo* transmission but not necessarily *in vitro*, depending on the cell line used (Davidson et al., 2020; Liu et al., 2020; Peacock et al., 2020; Sasaki et al., 2020). Structural data of SARS-CoV-2 spike indicates that cleavage at the S1/S2 boundary results in exposure of the receptor-binding domain (RBD) of spike (Wrobel et al., 2020). It has been suggested that this exposure of the RBD facilitates binding to ACE2 and the secondary cleavage of the S2’ site of spike, facilitating membrane fusion.

Interferons (IFNs) upregulate the expression of a range of antiviral proteins termed IFN-stimulated genes (ISGs) that inhibit various aspects of viral lifecycles, including entry (Schoggins, 2019). One of these protein families, IFN-induced transmembrane proteins (IFITMs) are membrane spanning proteins which inhibit the entry of several viruses, including HIV-1, influenza, Ebola and SARS-CoV-1 through blocking the fusion of the cellular and viral membranes, possibly by decreasing membrane fluidity or affecting membrane curvature (Foster, Pickering, & Neil, 2018; Zhao, Li, Winkler, An, & Guo, 2019). Three IFITMs demonstrate antiviral activity in humans: IFITM1 which localises to the plasma membrane, and IFITMs 2 and 3 which localise to late and early endosomes respectively (Foster et al., 2016; Shi, Schwartz, & Compton, 2017). Previous research has shown the route of entry correlates with the restriction of both influenza virus and HIV-1. Mislocalising IFITM3 to the cell surface abrogates IFITM3 restriction of influenza virus (Brass et al., 2009). In HIV-1, CCR5-tropic viruses which fuse at the plasma membrane are more restricted by IFITM1, and CXCR4-tropic viruses which utilise the endosomal route are more restricted by IFITMs 2 and 3 (Foster et al., 2016). It has been reported that SARS-CoV-2 is highly sensitive to type I and III IFNs, and more specifically, to IFITM3 (Felgenhauer et al., 2020; Lokugamage et al., 2020; Peacock et al., 2020; Stanifer et al., 2020). Conversely, other authors have suggested expression of IFITMs can enhance entry of SARS-CoV-2 (Bozzo et al., 2020). Given that entry is the first key step in viral transmission and IFITMs have been shown to be expressed in lung tissue, the interplay between IFITM restriction and SARS-CoV-2 entry is likely to be fundamental to the ability of SARS-CoV-2 to infect and transmit (Bailey, Zhong, Huang, & Farzan, 2014; Hachim et al., 2020). Here, we have shown the differential sensitivity of SARS-CoV-2 to IFITMs and how the presence of a polybasic cleavage site may affect entry in the context of IFITM restriction.

## RESULTS

### Sensitivity of SARS CoV-2 and pseudotyped lentiviral vectors (PLVs) with SARS-CoV-2 spike to human type-I, type II and type III interferons in A549-ACE2 cells

In order to examine the restriction of SARS CoV-2 replication by human antiviral proteins, we first sought to confirm the sensitivity of replication-competent SARS-CoV-2 (SARS-CoV-2 strain England 2) to type I (α and β), type II (γ) and type III (λ) IFNs in human A549 lung cancer cells stably expressing ACE2. We pre-treated the cells with different doses of recombinant human IFNα2, β, λ4 and γ overnight and then challenged them with SARS-CoV-2 (MOI 0.005 based on Vero-E6). 48 hours later we measured the levels of viral RNA by RT-qPCR using both Centres for Disease Control N1 and N2 primer probe sets (Figure 1A and D). We found that SARS-CoV-2 is highly sensitive to IFNβ and IFNγ with very low IC_50_ values, and less sensitive but nonetheless still restricted by IFNα and IFNλ. In addition to measuring intracellular viral RNA abundance in the IFN-treated cells, we infected Vero-E6 cells with the supernatant harvested from IFN-treated and infected A549-ACE2 cells and quantified the expression of nucleocapsid (N) protein 24 hours later. This assay measures the amount of infectious virus produced by the mock or IFN-treated cells and showed similar results (Figure 1B and E), thus confirming previous studies that the virus is highly IFN sensitive, particularly to IFNβ and IFNγ, indicating that a number of ISGs have direct antiviral effects against SARS-CoV-2.

**Figure 1.**
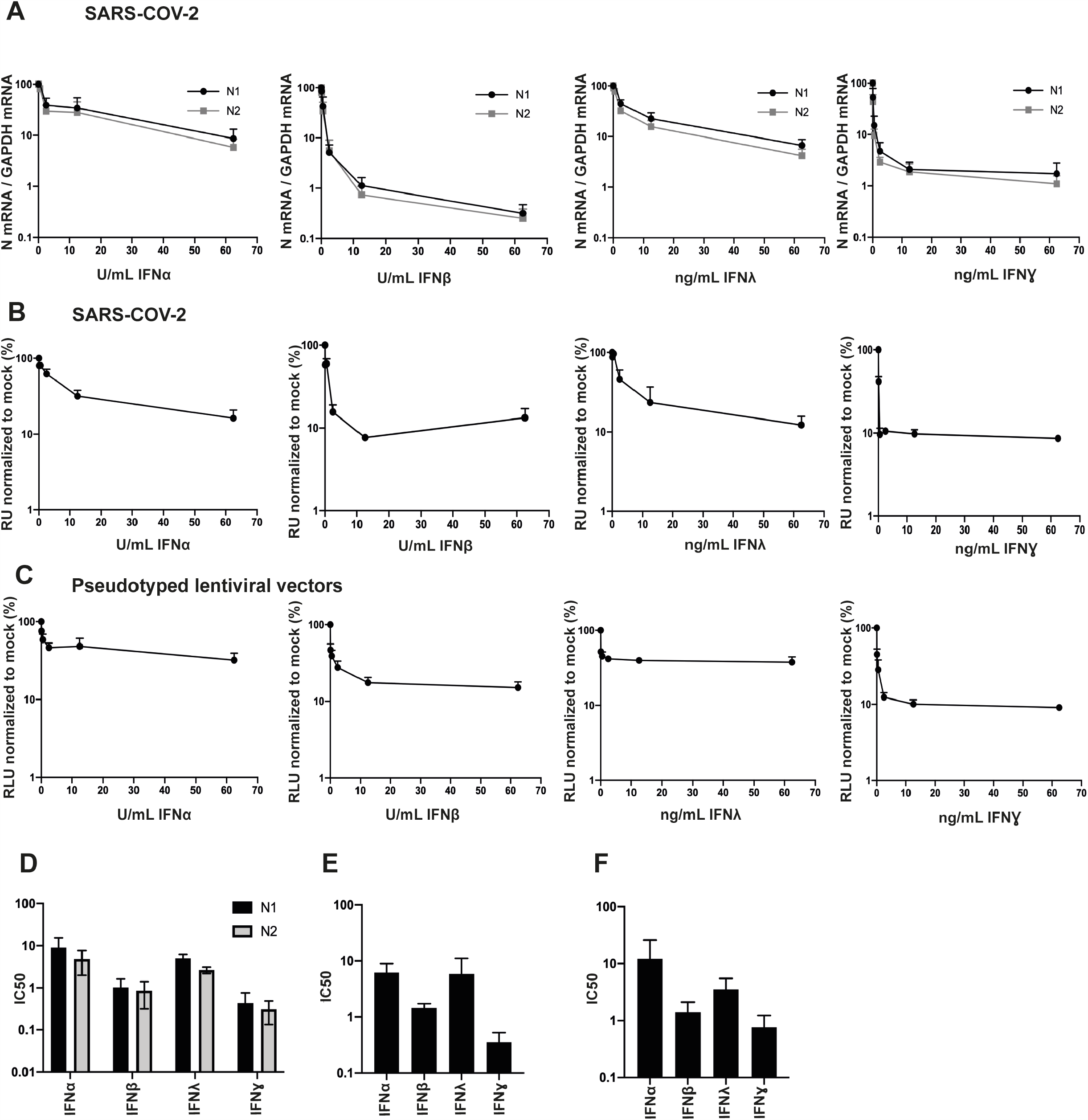
Entry of replication competent and PLVs of SARS-COV-2 is inhibited by IFNβ and IFN*γ*. **A)** A549-ACE2s were pre-treated for 18 hours with IFNα, β, λ, or ɣ and subsequently infected with replication-competent SARS-COV-2. Viral RNA was extracted 48 hours later and detected with two sets of primers (N1 and N2) against nucleocapsid mRNA and normalized to infection in mock-treated cells. N=3, Mean±SEM. **B)** Supernatant from infected A549-ACE2s in A) was used to infect Vero-E6 cells for 24 hours. Vero-E6s were then stained for nucleocapsid protein and normalized to mock-treated condition. RU = relative units. N=3, Mean±SEM. **C)** A549-ACE2s were pre-treated for 18 hours with IFNα, β, λ, or ɣ and transduced with PLVs of SARS-COV-2 for 48 hours. Infection was quantified by luciferase activity and normalized to mock-treated condition. RLU = relative luminescence units. N=3, Mean±SEM. **D)** IC50s of A) were calculated in Prism. N=3, Mean±SEM. **E)** IC50s of B) were calculated in Prism. N=3, Mean±SEM. **F)** IC50s of C) were calculated in Prism. N=3, Mean±SEM.

In order to address the activities of ISGs directed against spike-mediated entry, we first determined whether we could recapitulate the IFN phenotypes observed above using pseudotyped lentiviral vectors (PLVs). We generated PLVs containing SARS-CoV-2 spike bearing a luciferase reporter gene and tested them for sensitivity to IFNs on A549-ACE2. Similar to full-length SARS-CoV-2, we found that PLVs with SARS-CoV-2 spike are also highly sensitive to IFNβ and IFNγ (Figure 1C and F). Whilst the early events of HIV-1 are known targets of IFN treatment in some cell lines, these data suggest that when we isolate the entry stage of SARS-CoV-2 infection we observed inhibition by IFNβ and IFNγ (Doyle, Goujon, & Malim, 2015).

### SARS-CoV-2 is sensitive to IFITM2, but not IFITM3, in A549-ACE2 cells

IFITMs are a family of ISGs upregulated by IFNs that predominantly inhibit fusion of viral and cellular membranes (Foster et al., 2018; Shi et al., 2017). Considering that our PLVs with SARS-CoV-2 spike demonstrated a similar extent of inhibition by IFNs to the full-length virus we suspected that IFITMs, which have previously been reported to inhibit SARS-CoV-1 and more recently suggested to inhibit SARS-CoV-2, may be contributing to this inhibition (Bozzo et al., 2020; Huang et al., 2011; Peacock et al., 2020; Zhao et al., 2019).

To test the impact of each individual IFITM on SARS-CoV-2 infection, we generated stable A549-ACE2 cell lines expressing each human antiviral IFITM (Figure 2A) and infected these with PLVs of SARS-CoV-2. We found that SARS-CoV-2 showed a small but significant sensitivity to IFITM1 and a comparatively greater sensitivity to IFITM2 (Figure 2B). We recapitulated these phenotypes by challenging the A549-ACE2-IFITM cells with SARS-CoV-2 at increasing MOIs and using the supernatant of these cells 48 hours later to infect Vero-E6 cells, and measuring viral infectivity by staining for N protein (Figure 2C). At low MOI, SARS-CoV-2 was particularly sensitive to IFITM2 but not IFITM3, with an inhibitory effect seen with IFITM1, and these sensitivities were ameliorated at high viral inputs. As both single round PLVs and the full-length virus essentially displayed similar phenotypic sensitivity to IFITM2, these results suggest that a predominant antiviral effect is mediated at cellular entry.

**Figure 2.**
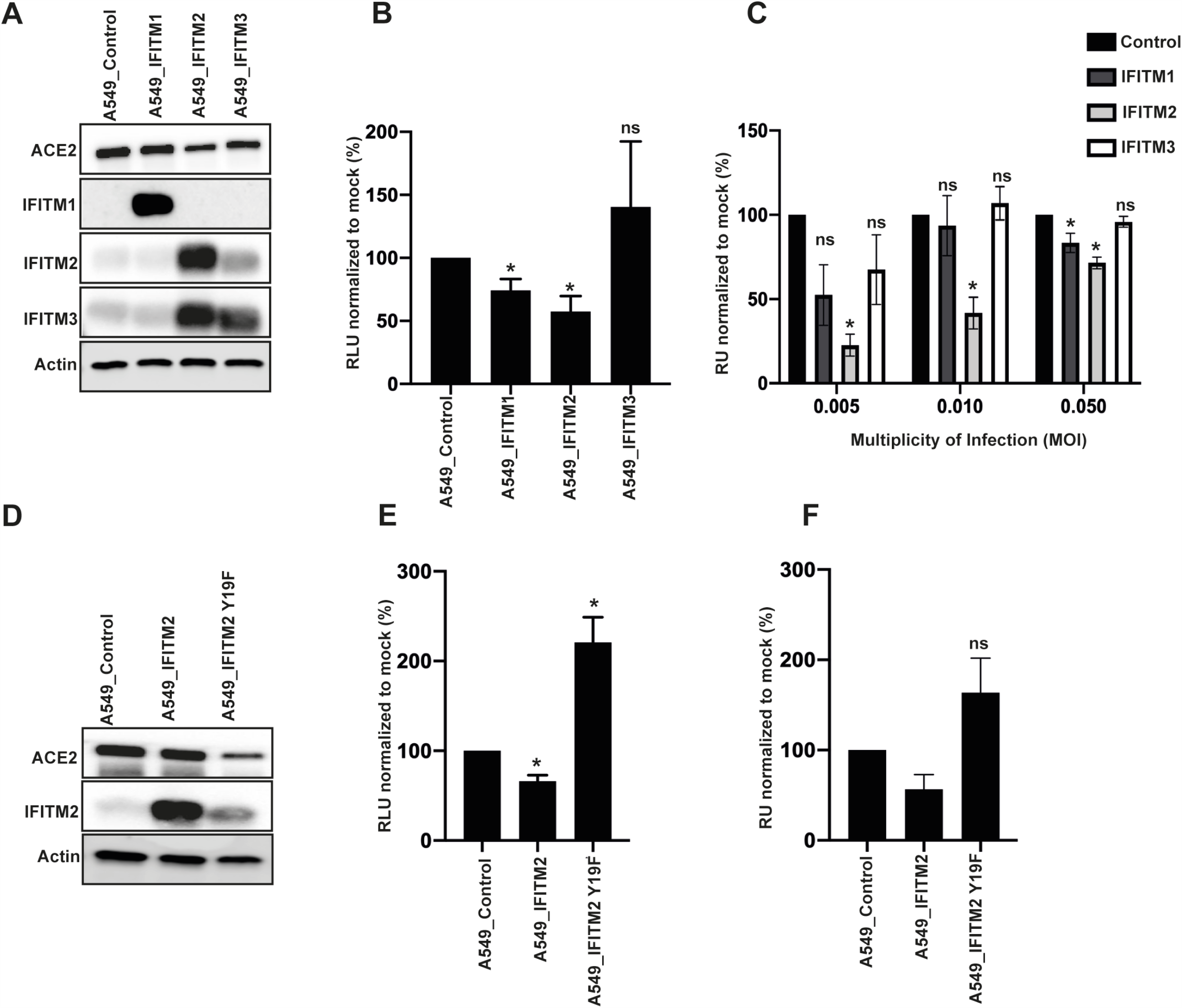
Replication competent and PLVs of SARS-COV-2 are inhibited by IFITM2 in A549-ACE2s. **A)** Representative immunoblot of A549-ACE2s stably expressing IFITM1, IFITM2, and IFITM3. Of note, the antibody to IFITM2 and IFITM3 recognises both proteins. **B)** A549-ACE2-IFITM cells were transduced with SARS-COV-2 PLVs for 48 hours and infection quantified by luciferase activity. RLU = relative luminescence units. N=3, Mean±SD. **C)** A549-ACE2-IFITM cells were infected with replication-competent SARS-CoV-2 for 48 hours at various MOIs. Supernatant was then used to infect Vero-E6 cells for 24 hours and stained for nucleocapsid protein. RU = relative units. N=3, Mean±SD. **D)** Representative immunoblot of A549-ACE2s stably expressing IFITM2 and IFITM2 Y19F. **E)** SARS-COV-2 PLVs were used to transduce A549-ACE2-IFITM2 and Y19F mutant and infection quantified 48 hours later by luciferase activity. RLU = relative luminescence units.N=3, Mean±SEM. **F)** Replication competent SARS-COV-2 was used to infect A549-ACE2-IFITM2 cells or A549-ACE2 cells stably expressing IFITM2 or IFITM2 Y19F at MOI 0.005. Supernatant was used to infect Vero-E6 cells for 24 hours and N stained as in C). Unpaired t test *=p=<0.05, N=3, Mean±SEM. RU = relative units.

Both IFITM2 and IFITM3 predominantly localize to endosomal compartments but reach them via endocytosis from the cell surface through the recruitment of the clathrin adaptor AP2 to a tyrosine-based endocytic signal (YxxΦ) in the IFITM2/3 cytoplasmic tail. We and others have previously demonstrated that mutating the Y19/Y20 to a phenylalanine in IFITM2 and IFITM3 respectively results in their accumulation at the plasma membrane (Chesarino, McMichael, Hach, & Yount, 2014; Foster et al., 2016). To test this, we stably expressed IFITM2 Y19F A549-ACE2 (Figure 2D) and infected these cells with both PLVs and replication competent SARS-CoV-2 and found that infection was not inhibited, but rather was enhanced by the presence of IFITM2 Y19F (Figure 2E, 2F). Although surprising that mislocalisation of IFITM2 resulted in enhancement of infection rather than simply an absence of restriction, these data are consistent with a recent report suggesting that similar mutants of IFITM3 enhance SARS-CoV-2 infection (Shi et al., 2020). These data suggest that the localisation of IFITM2 to endosomes or its recruitment to clathrin-coated pits at the plasma membrane is key to its inhibition of SARS-CoV-2 entry.

### The polybasic cleavage site determines sensitivity to IFITM2 in the presence or absence of TMPRSS2

A major difference between the spike protein of SARS-CoV-2 and the majority of the phylogenetically related bat Sarbecoviruses, including SARS-CoV-1, is the presence of the polybasic cleavage site at the S1/S2 boundary which facilitates the processing of spike to S1/S2 during viral assembly in the producer cell rather than during entry of the target (Figure 3A). As this feature has been proposed to be associated with the increased transmissibility of SARS-CoV-2, we hypothesized that it might affect the sensitivity of the virus to IFITM2. To investigate this, we deleted the polybasic cleavage site from SARS-CoV-2 (while preserving the adjacent RS cathepsin site) and swapped the corresponding region from (P_681_-A_684_) SARS-CoV-2 into SARS-CoV-1, generating SARS-CoV-2ΔPRRA and SARS-CoV-1 PRRA respectively (Figure 3A). We made PLVs of these mutants and analysed spike expression and virion incorporation by western blot using a polyclonal antibody against SARS-CoV-1/2 S2 (Figure 3B). We found that all spike proteins were equivalently expressed in the transfected producer 293T-17 cell. As expected, the SARS-CoV-1 spike exists predominantly as the S1/2 precursor on pelleted virions in the supernatant. By contrast, processed S2 was the predominant species found on virions pseudotyped with SARS-CoV-2 spike, indicating furin-mediated cleavage during virion assembly. By contrast, as expected SARS-CoV-2ΔPRRA was no longer cleaved. Insertion of the SARS-CoV-2 cleavage site into SARS-CoV-1 was sufficient to lead to processed spike, however this was not as efficient as in SARS-CoV-2, with virions incorporating both cleaved and uncleaved spikes (Figure 3B). In keeping with results from others, in Vero-E6 cells SARS-CoV-2ΔPRRA PLVs had a markedly *increased* infectivity of approximately 50-fold in A549-ACE2 cells compared to the wild type spike, approaching those of SARS-CoV-1 (Figure 3C) (Hoffmann, Kleine-Weber, & Pöhlmann, 2020). Addition of the PRRA site to SARS-CoV-1 slightly reduced titres. Since SARS-CoV-2 requires TMPRSS2 in the target cells to activate spike for entry, we over-expressed TMPRSS2 in A549-ACE2 cells by retroviral transduction and found that this specifically enhanced infection of SARS-CoV-2 PLVs, indicating that in the absence of TMPRSS2 expression, much of the SARS-CoV-2 inoculum is not infectious in these cells.

**Figure 3.**
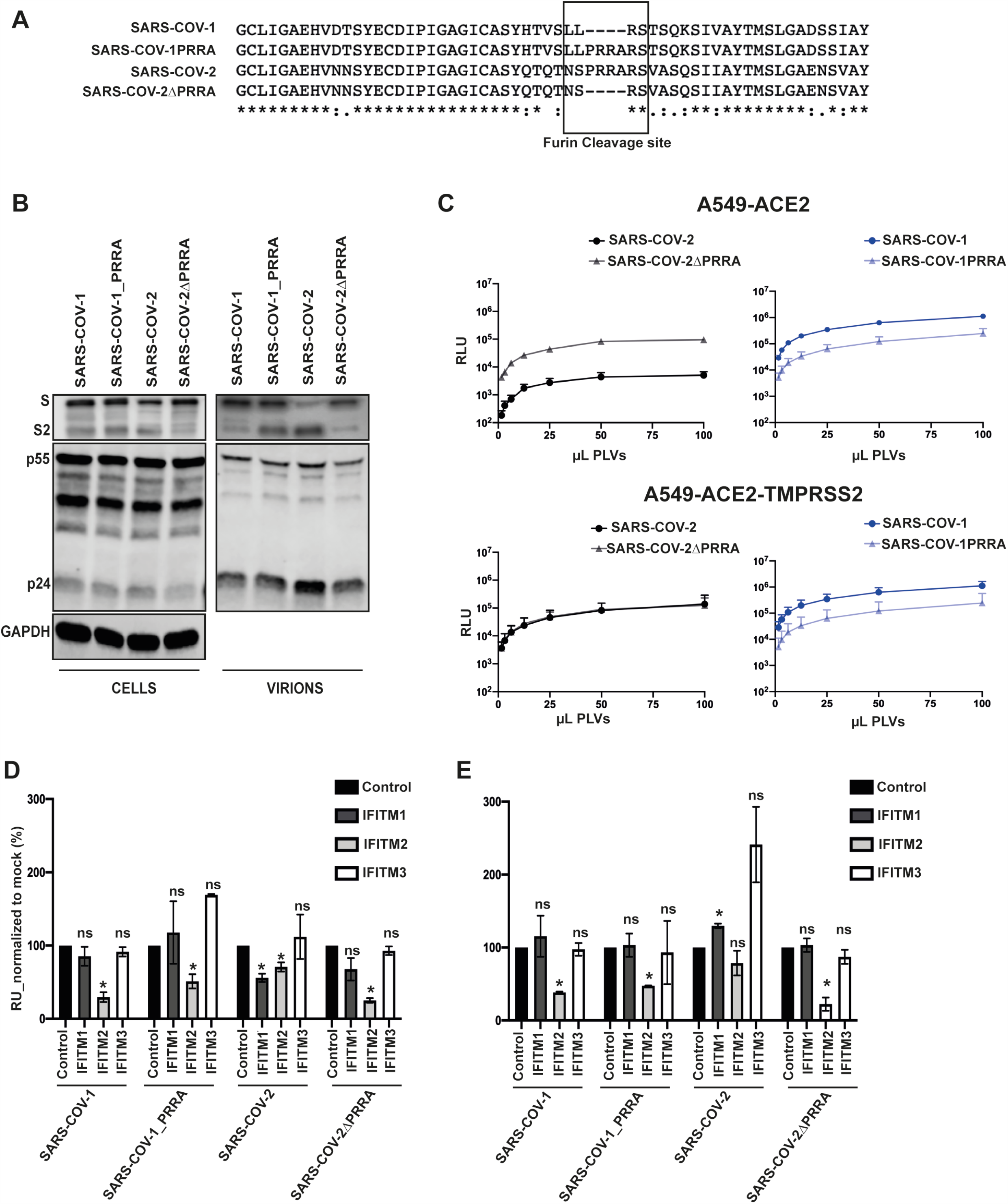
The presence or absence of a polybasic cleavage site determines sensitivity to IFITM2. **A)** Alignment of S1/S2 boundary in SARS-COV-1 and SARS-COV-2 with mutants where PRRA has been inserted/deleted. Alignment created in Clustal Omega. **B)** Representative immunoblot of PLVs of SARS-COV-1, SARS-COV-1_PRRA, SARS-COV-2, SARS-COV-2ΔPRRA. **C)** PLVs of SARS-COV-1, SARS-COV-1 PRRA, SARS-COV-2, and SARS-COV-2ΔPRRA were titrated on A549-ACE2 or A549-ACE2-TMPRSS2 cells and infectivity measured by luciferase 48 hours later. N=3, Mean±SEM. **D and E)** PLVs of SARS as in B) and C) were used to transduce A549-ACE2-IFITM cells for 48 hours and infection measured by luciferase activity. Infection was normalized to empty vector cells. Unpaired t test *=< p=0.05, RLU = relative luminescence units, N=3, Mean±SEM.

We then tested IFITM sensitivity of these PLVs in A549-ACE2 cells with and without TMPRSS2 overexpression (Figure 3D and 3E). As expected, SARS-CoV-2 was sensitive to both IFITM1 and IFITM2 in A549-ACE2 cells (Fig 3D). SARS-CoV-1 PLVs were significantly more sensitive to IFITM2 but displayed no restriction by IFITM1, suggestive of distinct subcellular site of entry between SARS-CoV-1 and SARS-CoV-2. Interestingly deletion of PRRA in SARS-CoV-2 rendered this spike as sensitive as SARS-CoV-1 to IFITM2 and slightly reduced the IFITM1 sensitivity. By contrast the addition of a cleavage site to SARS-CoV-1 significantly reduced IFITM2 sensitivity, albeit not to the levels of the fully cleaved SARS-CoV-2 spike. When we overexpressed TMPRSS2, we found that while IFITM1 sensitivity of SARS-CoV-2 could be abolished, this was not sufficient to rescue SARS-CoV-2 or SARS-CoV-2ΔPRRA from IFITM2. Thus, the presence of the polybasic cleavage site markedly reduces the sensitivity of SARS CoV-2 S-mediated entry to IFITM2, suggesting it affects the route of cellular entry into the cell and distinguishes it from SARS CoV-1.

To address the effects of spike cleavage on route of entry, we first determined the pH-sensitivity of the spike cleavage mutants using concanamycin A (ConA), an inhibitor of the vacuolar ATPase in late endosomes (Figure 4A). As expected, SARS-CoV-1 PLVs were exquisitely sensitive (1000 fold) to ConA inhibition in A549-ACE2 indicating that entry was occurring exclusively in a low pH endosomal compartment. In the presence of TMPRSS2, SARS-CoV-1 pH sensitivity was reduced, but entry still remained 20-50 fold lower suggesting any enhanced S2’ processing was not sufficient to abolish pH-dependent entry. Similarly, while insertion of a partially processed polybasic cleavage site in SARS-CoV-1 reduced but did not abolish pH-dependent entry in either cell type. By contrast, entry of SARS CoV-2 PLVs was only mildly affected (2-3 fold) by ConA treatment irrespective of TMPRSS2 overexpression, indicating that most viral entry was occurring at neutral pH, and TMPRSS2 was enhancing entry at this point rather than elsewhere in the cell. Similar to SARS-CoV-1, deletion of the PRRA site from SARS-CoV-2 rendered PLVs strictly pH-dependent without affecting titre. In keeping with these data, unlike SARS-CoV-2, SARS-CoV-1 and SARS-CoV-2ΔPRRA-mediated entry was inhibited by the endosomal cathepsin inhibitor E64D but not the TMPRSS inhibitor Camostat (Figure 4B-I). By contrast, SARS-CoV-2 only was sensitive to E64D in TMPRSS2-overexpressing cells. Together these data suggest that S1/S2 cleavage by furin in the producer cell promotes TMPRSS2-mediated entry at the plasma membrane or soon after internalization, and abolishes the requirement for cathepsin-mediated processing in the acidic endosomal compartments. The data further suggest that in the absence of abundant TMPRSS2 at the cell surface, the processed SARS-CoV-2 cannot efficiently enter through a low pH compartment. Thus, the PRRA site dictates the route of entry into the cell and thus its sensitivity to IFITM proteins that occupy different cellular locations.

**Figure 4.**
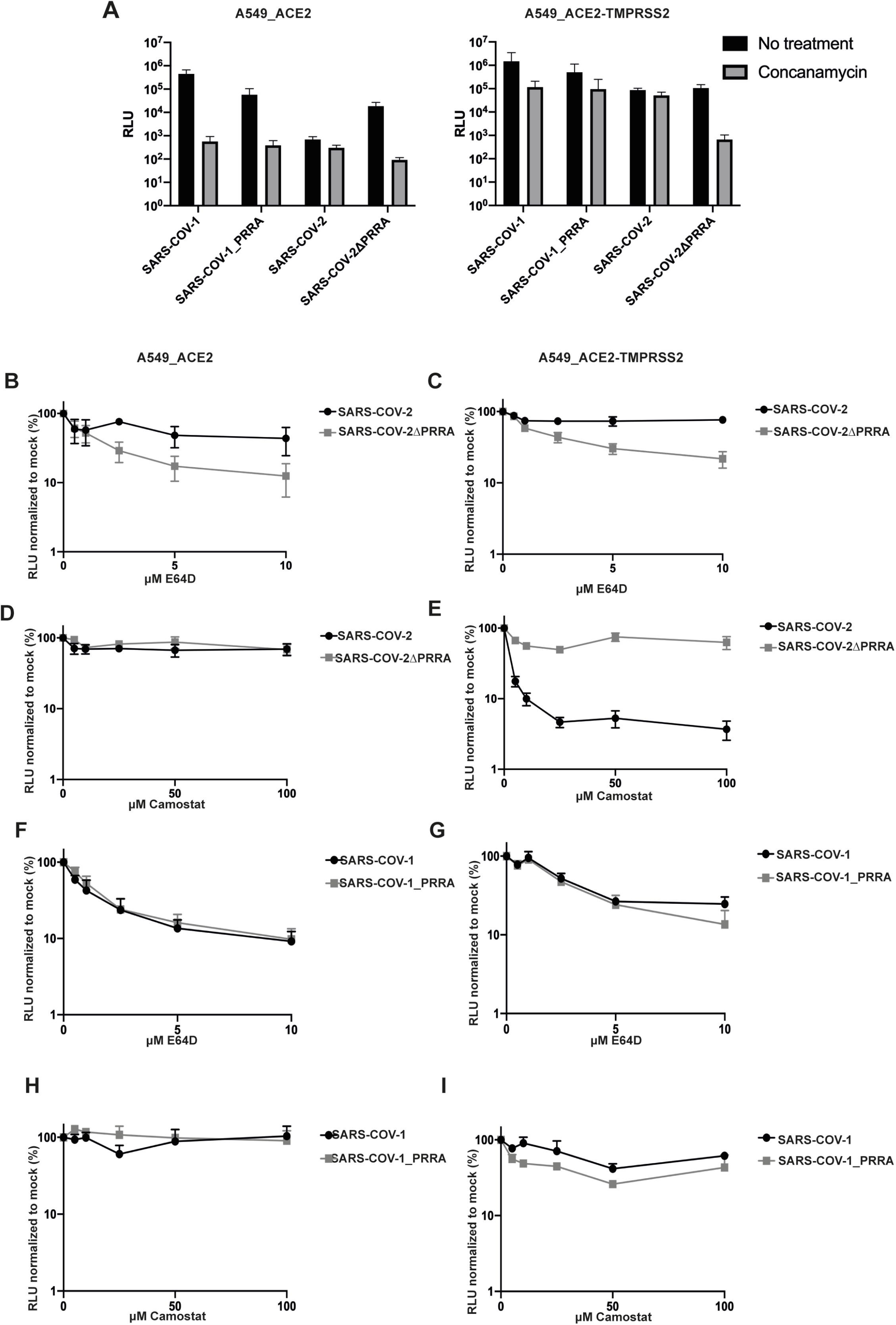
SARS-COV-2 and SARS-COV-2ΔPRRA differ in preferential route of entry. **A)** A549-ACE2 and A549-ACE2-TMPRSS2 cells were treated with 100nM of concanamycin for 1 hour and transduced with SARS PLVs. Infection was determined by luciferase activity 48 hours later. Black = non-treated, grey = concanamycin. RLU = relative luminescence units. **B, C, D, E)** A549-ACE2 or A549-ACE2-TMPRSS2 were treated for 1 hour with E64d or Camostat and subsequently transduced with PLVs of SARS-COV-2 or SARS-COV-2ΔPRRA and infection determined by luciferase activity 48 hours later. RLU = relative luminescence units. N=3, Mean±SEM. **F, G, H, I)** A549-ACE2 or A549-ACE2-TMPRSS2 were treated with E64d or Camostat for 1 hour and transduced with PLVs of SARS-COV-1 or SARS-COV-1 PRRA and infection determined by luciferase activity 48 hours later. RLU = relative luminescence units. N=3, Mean±SEM.

### IFITM2 contributes to the antiviral restriction of SARS-CoV-2 by IFNβ

Having established that IFITM2 can restrict SARS-CoV-2 depending on its mechanism of entry, we wanted to determine how much of the inhibition of replication-competent SARS-CoV-2 by IFNβ and IFNγ could be attributed to IFITM2. We examined the expression of IFITM2 and IFITM3 in IFN-treated A549-ACE2 and observed a robust upregulation of both IFITM2 and IFITM3 following treatment with IFNβ. By contrast, while IFITM3 was also robustly induced by IFNγ, IFITM2 was weakly induced (Figure 5A). Using siRNAs against IFITM2 that rescued SARS-CoV-2 replication in A549-ACE2-IFITM2 cells (Figure 5B, 5C), we then knocked down IFITM2 in the context of pre-treating A549-ACE2 cells with IFNβ or IFNγ and challenged the cells with SARS-CoV-2, measuring infectious virus output on Vero-E6 cells 48h later (Figure 5D and E). IFITM2 depletion substantially relieved the inhibition of viral replication by IFNβ treatment, whereas that induced by IFNγ was only modestly alleviated. This was reflected in the 20-fold increase in the IC_50_ of IFNβ, but only a 2-fold increase in IFNγ, indicating that in these cells IFITM2 is a major component of the type I IFN-antiviral state protecting cells from SARS-CoV-2 (Figure 5F). Furthermore, in A549-ACE2 cells overexpressing TMPRSS2, the knockdown of IFITM2 essentially abolished all the antiviral activity of pretreating the cells with IFNβ (Figure 5G). Thus IFITM2-mediated entry restriction is a major type-I IFN activity that constitutes an antiviral state, blocking the replication of SARS CoV-2.

**Figure 5.**
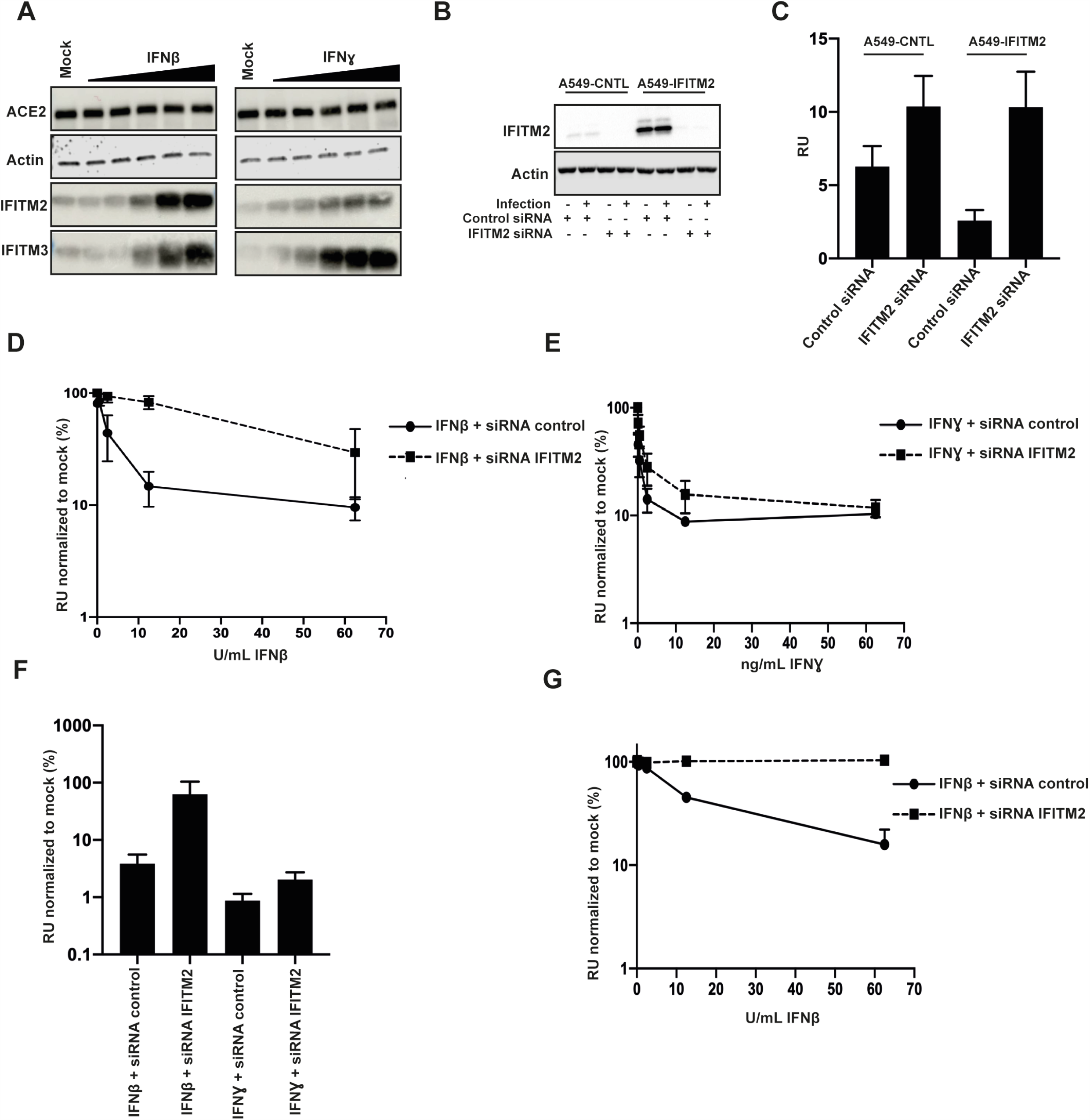
SiRNA of IFITM2 rescues IFNB mediated restriction of replication competent SARS-COV-2. **A)** Representative immunoblot of A549-ACE2 treated with different amounts of IFNβ or IFNγ for 18 hours. **B, C)** A549-ACE2s were transfected with siRNAs against non-targeting control or IFITM2 and supernatants used to infect Vero-E6 cells for 24 hours and stained for nucleocapsid protein. RU = relative units. N=3, Mean±SEM. D) A549-ACE2 were pre-treated with IFNβ and IFNɣ for 18 hours and infected with replication-competent SARS-COV-2. Infected supernatant was used to infect Vero-E6 cells for 24 hours and cells stained for N protein. N=3, Mean±SEM. **F)** IC50s of D) and E) were calculated in Prism. N=3, Mean±SEM. **G)** A549-ACE2-TMPRSS2 were transfected with siRNAs against non-targeting control or IFITM2 when seeding and prior to IFN treatment. Cells were treated with IFNβ and infected with replication competent SARS-COV-2 18 hours later. Infected supernatant was used to infect Vero-E6 cells for 24 hours and cells stained for N protein. N=3, RU = relative units. Mean±SEM.

## DISCUSSION

In this study we have provided evidence that IFITM2 has potent inhibitory activity against SARS-CoV-2 entry and constitutes at least part of the antiviral activity conferred by treatment of target cells with IFNβ. Furthermore, we find that the presence or absence of the polybasic cleavage site, which facilitates pH-independent entry of SARS-CoV-2, modulates the sensitivity of the virus to IFITM2

In contrast to SARS-CoV-1 and other related SARS-like CoVs in bats, SARS-CoV-2 is distinguished by the presence of a furin cleavage site at the S1/S2 boundary. This leads to the spike on SARS-CoV-2 virions being predominantly cleaved in the producer cell rather than by cathepsins during endocytic entry into the target cell. This renders its entry pH-independent, suggesting fusion occurs at, or near, the cell surface. Recent evidence further indicates that the furin cleavage generates a C-terminal ligand on S1 that interacts with neuropilin-1 (NRP-1) on the surface of target cells in the lung (Cantuti-Castelvetri et al., 2020; Daly et al., 2020). The role of NRP-1 is not completely clear, but there is some suggestion that it may stabilize the attachment of SARS-CoV-2 at the cell surface to facilitate either ACE2 interaction or processing of the S2’ site by TMPRSS2. Structural analyses of the SARS-CoV-2 spike trimer further shows that furin-mediated cleavage facilitates at least one RBD domain to adopt an erect conformation that would further promote ACE2 interaction (Wrobel et al., 2020). Interestingly, deletion of the PRRA is not detrimental to SARS-CoV-2 entry in all cell types in culture, and in fact in TMPRSS2-low Vero-E6 cells the furin-cleavage site is rapidly lost upon passage, suggesting it can actively hinder infection. Herein we show that while wild-type spike-mediated entry is insensitive to inhibition of endosomal pH, the cleavage mutant is strictly dependent on endosome acidification and cathepsins. Interestingly, for efficient entry, SARS-CoV-2 requires high TMPRSS2 expression to activate the fusion mechanism by cleaving S2’. However, in cells where TMPRSS2 is limiting, SARS-CoV-1 and SARS-CoV-2ΔPRRA entry is far more efficient. Thus, in its uncleaved form SARS-CoV-2 spike can mediate entry in endosomes, but in its mature form entry cannot be rescued in low pH compartments of TMPRSS2-low cells. This would imply that the cleaved spike is unstable at endosomal pH, and interestingly recent studies from the Kwong group indicate that conformational dynamics of the RBD are also pH sensitive (Zhou et al., 2020). Despite this potential greater fragility of the SARS-CoV-2 trimer, the furin cleavage site appears to be essential for replication in primary airway epithelium and for transmission in ferret models (Peacock et al., 2020). We suggest that one of the reasons pH-independent fusion at or near the cell surface is maintained is to mitigate the antiviral activity of IFITM proteins, particularly IFITM2.

The localization of IFITMs largely defines which viruses they can restrict. Whilst they can be incorporated into nascent virion membranes and exert an antiviral effect there, their best studied mechanism of action is to prevent fusion of an incoming virus at the target cell membrane. IFITM1 is predominantly found at the plasma membrane, whereas IFITM2 and IFITM3 occupy endosomal compartments by virtue of a conserved endocytic signal. Palmitoylation of the intracellular loop of the IFITM stabilizes their conformation in the membrane and promotes their homo- and heterotypic interactions. The current model for their action is that IFITM:IFITM interactions exert a level of positive curvature to the target membrane that arrests enveloped viral entry at the hemi-fusion stage (Rahman et al., 2020). IFITM3 is particularly potent against influenza viruses and its redistribution away from early endosomes by mutating the endocytic site in the cytoplasmic tail abolishes its antiviral activity. Comparatively less is known about IFITM2, although it has been shown to inhibit a number of other enveloped viruses that enter in later endosomes. Of note, human IFITM2 and 3 differ from each other by only 10 amino acids, and yet their restriction patterns are not interchangeable. While IFITM2 and 3 are localized in endosomal compartments, they traffic via the cell surface, and their recruitment into clathrin coated pits would imply that they may have some activity at viral entry sites at the plasma membrane as well. However, our observations that IFITM2, but not IFITM3, in the A549 system inhibits SARS-CoV-1 and CoV-2 would suggest that neither virus fuse significantly in a cellular compartment occupied by IFITM3. Cell type specificity in IFITM localization (due to endocytic rate etc) and hetero-typic interactions between IFITMs suggests that when all are co-expressed, they may form a more complex barrier to enveloped virus fusion than an individual IFITM alone.

Studies on SARS-CoV-1 and recent papers and preprints on SARS-CoV-2 have shown a variety of phenotypes with different IFITMs on both viral entry and cell-to-cell fusion mediated by the spike protein(Bozzo et al., 2020; Peacock et al., 2020; Shi et al., 2020). IFITM1 appears to block syncytium formation between infected and uninfected cells and this is overcome by TMPRSS2 expression, which is in keeping with our observations that in stably expressing cells small effects of IFITM1 on SARS-CoV-2 entry in A549-ACE2 cells can be abolished similarly (Buchrieser et al., 2020). Other data has implicated IFITM3, and demonstrated that it can be enhancing if its expression is restricted to the cell surface. Most of these experiments have been performed in transiently transfected 293T cells with PLVs, and whilst the known determinants of IFITM3 function are required, whether transient overexpression faithfully represents the localization and potency of IFITM2 and IFITM3 natural expression is unclear. Here we find that stable ectopic expression of IFITM2 and to some extent IFITM1 restricts both the entry of PLVs and the replication of the SARS-CoV-2 virus itself in A549-ACE2 cells. The enhanced sensitivity of the PRRA mutant of SARS-CoV-2 and SARS-CoV-1 to IFITM2 is entirely consistent with their dependence on low pH compartments for cathepsin cleavage. By restricting IFITM2 to the plasma membrane and outside of clathrin coated pits by abolishing AP2 interaction, we see similar enhancement effects to those seen by the Yount group with IFITM3 (Shi et al., 2020). Why this happens is not known, but given the effects that IFITMs have on membrane fluidity, this may be an indirect effect on the surface levels and distribution of entry cofactors at the plasma membrane. It also suggests why there may be an association of the rs12252-C polymorphism that expresses an N-terminally truncated IFITM3 with COVID-19 severity (Gómez et al., 2021). Restriction by IFITM2 but not IFITM3 is surprising. This would suggest that IFITM2 localization is not only limited to later endosomes than IFITM3, but may also reside in distinct localizations at or near the plasma membrane dependent on its AP2-binding site.

In addition to examining the sensitivity of SARS-CoV-2 to individual IFITM proteins, we also showed that IFITM2 knockdown is sufficient to alleviate much of the antiviral effect of pretreating A549 cells with type I, but not type II IFN. Studies from many groups have shown that while SARS-CoV-2 is a poor inducer of IFN-responses in infected cells early in the replication cycle, it is highly sensitive to pretreatment of target cells by both type I, II and III IFNs (Lokugamage et al., 2020; Nchioua et al., 2020; Stanifer et al., 2020). This suggests the potential for multiple ISGs to restrict SARS-CoV-2 replication and has raised the possibility of IFNs as possible treatments for COVID-19 (Haji Abdolvahab et al., 2020). The role of IFNs in SARS-CoV-2 pathogenesis is complex. Genetic lesions in pattern-recognition and IFN signaling as well as serum autoantibodies that neutralize type I IFNs are associated with risk of severe COVID-19(Bastard et al., 2020). However, dysregulated or delayed IFN responses driving systemic inflammation may underlie some of the pathology in COVID-19 (Lucas et al., 2020). Understanding which aspects of the IFN-response are antiviral against SARS CoV-2 is thus of very high importance.

In A549-ACE2 cells, IFITM2 is more potently induced by IFNβ than IFNγ and its knockdown substantially reduces the sensitivity of the virus to IFNβ-induced restriction. The sequence similarity between IFITM2 and IFITM3 means that it is difficult to knockdown one without affecting the other. The lack of IFITM3 restriction when expressed alone, and its potent expression after both IFNβ and IFNγ treatment would argue against IFITM3 playing the major role. However, given that IFITMs can interact with each other, we cannot rule out that IFITM1 or IFITM3 play a role in potentiating IFITM2’s antiviral activity after IFN induction. The former is a distinct possibility as IFITM2 knockdown fully rescues SARS-CoV-2 from IFNβ treatment in cells over-expressing TMPRSS2. Since we found that the minor restriction conferred by IFITM1 alone is abolished by TMPRSS2 expression, a plausible explanation is that more robust S2’ activation of SARS-CoV-2 spike at the cell surface overcomes IFITM1inhibition by saturating its activity (Weston et al., 2016). While it is surprising that IFNβ has no effect in these cells when IFITM2 is knocked-down, we would caution against interpreting that IFITM2 is the only ISG targeting SARS-CoV-2 replication. The rapidity and burst-size of SARS-CoV-2 replication in culture may render other relevant antiviral proteins difficult to measure. Furthermore, the virus encodes a number of antagonists of antiviral pathways (Xia et al., 2020). As shown clearly by the IFNγ phenotype, expression of other ISGs or their differential regulation may make a give antiviral more less potent. Of note, the IFNγ-mediated inhibition of SARS-CoV-2 is in part mediated through the zinc-finger antiviral protein (ZAP) (Nchioua et al., 2020).

Despite the sensitivity of SARS-CoV-2 to IFITM2, deletion of the PRRA cleavage site in spike substantially potentiates its antiviral activity. In most cells in culture expressing low levels of TMPRSS2, furin cleavage is detrimental to entry, and in A549 cells this can be rescued to the mutant levels of entry by ectopic expression of TMPRSS2. In primary lung epithelial cells, however, the wildtype spike is clearly superior and out competes the mutant as well as being more transmissible in ferret models (Peacock et al., 2020). Epithelial barrier tissues constitutively express a level of ISGs through the tonic activity of type I IFNs (Broggi et al., 2020). Therefore, it is tempting to speculate that the selection pressure for maintaining this attribute in SARS-CoV-2 spike is in part to promote cell surface fusion in target cells that already express IFITM2. Interestingly, Peacock et al have shown that an equivalent PRRA mutant virus can be rescued in lung epithelial cells by the antifungal drug amphotericin B, known to disrupt IFITM function (Peacock et al., 2020). Addition of a partially active PRRA cleavage site is not sufficient to reduce the IFITM2 restriction of SARS-CoV-1 S to that of SARS-CoV-2, and thus other determinants in spike are likely to modulate sensitivity.

In sum we show that IFITM2 is a key antiviral protein targeting SARS-CoV-2 entry, and its activity is modulated by the furin-cleavage site in spike. These data therefore suggest that therapeutic strategies which upregulate IFITM2 in epithelial tissues or inhibit furin-mediated cleavage of spike may render the virus more sensitive to innate-immune mediated control.

## METHODS

### Cell lines and plasmids

293T-17 (ATCC), A549-ACE2, A549-ACE2-TMPRSS2, Calu3 (ATCC), and A549-ACE2 expressing the individual IFITM proteins were cultured in DMEM (Gibco) with 10% FBS (Invitrogen) and 200ug/ml Gentamicin (Sigma). Codon optimised SARS-CoV-1 spike was synthesised by GeneArt and codon optimised SARS-CoV-2 spike and ACE2 were kindly given by Dr Nigel Temperton. Plasmid containing TMPRSS2 gene was kindly given by Dr Caroline Goujon. Mutants of spikes or IFITMs were generated with Q5® Site-Directed Mutagenesis Kit (E0554) following the manufacturers instructions: SARS-CoV-2 spike ΔPRRA (AGAAGCGTGGCCAGCCAG, GCTATTGGTCTGGGTCTGGTAG), SARS-CoV-1 spike PRRA (AGAGCCCGGAGCACCAGCCAGAAA, TCTAGGCAGCAGAGACACGGTGTG), IFITM2 Y19A (GCCTCCCAACgctGAGATGCTCAAGGAGGAG, TGGCCGCTGTTGACAGGA), IFITM2 Y19F (GCCTCCCAACtttGAGATGCTCAAGGAG, TGGCCGCTGTTGACAGGA)

A549 stable cell lines expressing ACE2 (pMIGR1-puro), TMPRSS2 (IRES-neo.WPRE, and IFITMs (pLHCX) were generated through transducing cells with lentiviral vectors packaged with HIV gag-pol (8.91) or MLV gag-pol and VSV-G. Cells were incubated with lentiviral vectors for 4 hours. Corresponding antibiotic selection was added 24 hours after transduction.

### Production of Pseudotyped Lentiviral Vectors and infection

293T-17 were transfected with firefly luciferase expressing vector (CSXW), HIV gag-pol (8.91) and spike at a ratio of 1.5:1:0.9ug using 35ul of PEI-MAX as previously described (Grehan, Ferrara, & Temperton, 2015). Media was changed 18 hours later and vectors were harvested through a 0.45um filter 48 hours after transfection. Viral supernatant was then used to transduce each cell line of interest for 48 hours and readout measured by Promega Steady-Glo® (E2550).

### Passage of SARS-CoV-2

PHE England strain 2 was propagated in Vero-E6 cells and titred by standard method.

### Infection with replication competent SARS-CoV-2

1.5×10^5^ A549-ACE2 cells were infected for 1h at 37°C with SARS-CoV-2 replication competent with an MOI of 0.005. Media was replaced and cells were incubated for 48h at 37°C. 48h later, cells were harvest for RNA extraction or protein analysis and the supernatant was used to infect Vero-E6 cells to measure virus infectivity.

### Interferon assays

Cells were treated with different doses of IFNα (Invitrogen, 111001) IFNβ (PBL Assay Science, 11415-1), IFNγ (Peprotech, 300-02) or IFNλ (Peprotech, 300-02L) for 18 hours prior to infection. Media was changed for virus or PLVs the following day and the infection was measured by Steady-Glo, qPCR or N-staining 48 hours later.

### siRNA knockdown of IFITM2

1×10^5^ A549-ACE2 cells were reverse transfected using 20pmol of Non-targeting siRNA(D-001206-13-20) or IFITM2 siRNA (M-020103-02-0010) and 1μL of RNAi max (Invitrogen). Cells were incubated for 24h prior a second round of reverse transfection. 8h later, cells were treated with different doses of IFNβ or IFNγ as previously described.

Following 16h of IFN treatment cells were infected with replication competent SARS-CoV-2 with an MOI of 0.005, as previously described. 48h after infection, cells were harvested for protein analysis and the supernatant was used to measure virus infectivity by N-staining.

### RT-qPCR

RNA from infected cells was extracted using QIAGEN RNeasy (QIAGEN RNeasy Mini Kit, 74106) following the manufacturers instructions. 1μL of each extracted RNA was used to performed one step RT-qPCR using TaqMan Fast Virus 1-Step Master Mix (Invitrogen). The relative quantities of nucleocapsid (N) gene were measured using SARS-CoV-2 (2019-nCoV) CDC qPCR Probe Assay (IDT DNA technologies).

### SARS-CoV-2 N-staining

2×10^4^ Vero-E6 cells were infected for 1h with 50μL of undiluted or 1/10 dilution of virus supernatant. Following infection 50μL of 2x overlay (DMEM + 2% FBS + 0.1% agarose) media was added to infected cells. 24h later, cells were fixed using 4% paraformaldehyde during 30 min at room temperature. Fixed cells were permeabilised with 0.1% Triton for 15 minutes, then blocked using 3% milk and incubated for 45 min with anti-human anti-SARS-CoV-2 N (CR3009). Following incubation, cells were washed with 1X PBS and incubated with secondary antibody, goat anti-human IgG (Fc) peroxidase conjugate (Sigma A0170) for 45 min. Finally, the presence of N protein was determined using 1-Step™ Ultra TMB-ELISA Substrate Solution (ThermoFisher).

### Drug assays

Cells were pre-treated with Camostat mesylate (Sigma, SML0057), E64D (Sigma, E8640), or Concanamycin (Cayman chemicals, 80890-47-7) for 1 hour at 37°C prior transduction. Cells were then transduced with PLVs for 48 hours and infection determined by luciferase activity.

### SDS-PAGE and Western blotting

Cellular samples were lysed in reducing Laemmeli buffer at 95°C for 10 minutes. Supernatant samples were centrifuged at 18,000 RCF through a 20% sucrose cushion for 1 hour at 4°C prior to lysis in reducing Laemelli buffer. Samples were separated on 8–16 % Mini-PROTEAN® TGX™ Precast gels (Bio-Rad) and transferred onto nitrocellulose membrane. Membranes were blocked in milk prior to detection with specific antibodies: 1:1000 ACE2 rabbit (Abcam, Ab108209), 1:1000 TMPRSS2 rabbit (Abcam, Ab92323), 1:2000 Actin mouse (Abcam, Ab6276), 1:5000 GAPDH rabbit (Abcam, Ab9485), 1:5000 HSP90 mouse (Genetex, Gt×109753), 1:50 HIV-1 p24Gag mouse (Chesebro, Wehrly, Nishio, & Perryman, 1992), 1:1000 Spike mouse (Genetex, Gtx632604), 1:3000 IFITM1 mouse (Proteintech, 60074-1-Ig), 1:3000 IFITM2 rabbit (Proteintech, 12769-1AP), 1:3000 IFITM3 rabbit (Proteintech, 11714-1-AP). Proteins were detected using LI-COR and ImageQuant LAS 4000 cameras.

## ACKNOWLEDGEMENTS

We are grateful to Nigel Temperton, Caroline Goujon, Andrew Davidson and Public Health England for reagents. We thank Michael Malim, Rui Pedro Galao, and Jose Jimenez-Guardeno for helpful advice. Finally, we thank all other members of the Neil and Swanson groups for help and good humour throughout a very difficult year.

## AUTHORS CONTRIBUTION

All experiments were performed by HW, MJL and AR. SP provided help in setting up the SARS-CoV-2 N immunostaining. KJD and CMS provided reagents, funding support and advice. HW, MJLB and SJDN analyzed the data and wrote the manuscript. All authors edited the manuscript and provided comments.

